# A Computational Dissection of Spike protein of SARS-CoV-2 Omicron Variant

**DOI:** 10.1101/2021.12.17.473260

**Authors:** Zaira Rehman, Massab Umair, Aamer Ikram, Muhammad Salman, Syed Adnan Haider, Muhammad Ammar

## Abstract

The emergence of SARS-CoV-2 omicron variant in late November, 2021 and its rapid spread to different countries, warns the health authorities to take initiative to work on containing its spread. The omicron SARS-CoV-2 variant is unusual from the other variants of concerns reported earlier as it harbors many novel mutations in its genome particularly with >30 mutations in the spike glycoprotein alone. The current study investigated the variation in binding mechanism which it carries compared to the wild type. The study also explored the interaction profile of spike-omicron with human ACE2 receptor. The structure of omicron spike glycoprotein was determined though homology modeling. The interaction analysis was performed through docking using HADDOCK followed by binding affinity calculation. Finally, the comparison of interactions were performed among spike-ACE2 complex of wild type, delta and omicron variants. The interaction analysis has revealed the involvement of highly charged and polar residues (H505, Arg498, Ser446, Arg493, and Tyr501) in the interactions. The important novel interactions in the spike-ACE2-omicron complex was observed as S494:H34, S496:D38, R498:Y41, Y501:K353, and H505:R393 and R493:D38. Moreover, the binding affinity of spike-ACE2-omicron complex (−17.6Kcal/mol) is much higher than wild type-ACE2 (−13.2Kcal/mol) and delta-ACE2 complex (−13.3Kcal/mol). These results indicate that the involvement of polar and charged residues in the interactions with ACE2 may have an impact on increased transmissibility of omicron variant.

## Introduction

The SARS-CoV-2 B1.1.529 Omicron variant initially emerged in Botswana and then subsequently reported in South Africa on 24^th^ November 2021. On 26 November 2021, the WHO listed the Omicron as Variants of Concern (VOC). The variant has been consistently in the headlines since then owing to its high transmissibility potential[1]. Till date the Omicron variant has spread to 42 countries. Within South Africa, it has been presumed that the number of affected cases has been doubled in less than 3 days, with 70 % of new SARS-CoV-2 cases turning out to be of omicron variant. The Omicron variant harbors more than 30 spike protein changes many of which have also been found in alpha and delta variants of SARS-CoV-2 [2]. There have been 26 unique mutations identified in the spike protein of omicron. In the already reported VOCs, no insertion have been observed in the spike protein, but omicron harbors a unique insertion ins214EPE. The detailed analysis of this insertion has shown that the same rearrangement have also been present in the spike protein of HCoV-229E. The possible explanation of this insertion may have been attributed to the person being co-infected with both the viruses and genetic recombination event resulted in the ins214EPE in omicron spike protein. The omicron variations may also have been conferring the variant with immune evasive potential, thus also raising the possibility of reducing effectiveness of SARS-CoV-2 vaccines. This notion gets stronger considering a quarter of South African population have been fully vaccinated. Recently Schmidt et al. 2021 has engineered a ‘polymutant’ spike capable of blunting the effects of neutralizing antibodies. Many of the mutations investigated by Schmidt et al has also been found in the omicron variant [3]. Within almost one month since the detection of variant a new version of Omicron has been identified and referred as BA.2 nicknamed as “Stealth Omicron”.

Due to large number of novel mutations in the receptor binding domain of omicron variant, it is important to study how this new variant interact with the ACE2 receptor. In order to understand the interaction pattern of omicron spike glycoprotein with ACE2, current study has been conducted.

## Materials and Methods

### Sequence Retrieval

The representative sequences of delta variant and omicron variant were retrieved from GISAID (GISAID ID: EPI_ISL_6649016 and EPI_ISL_2757739). The reference sequence of SARS-CoV-2 WHU01 was retrieved from NCBI (MN988668). The multiple sequence alignment was performed using CLUSTALW. The mutation profile of spike glycoprotein in delta and omicron variant was verified by sequence alignment with WHU01 sequence.

### Structure prediction and interaction analysis

The structure of spike glycoprotein complex with human ACE2 was retrieved from protein data bank (pdb ID: 7DF4) (referred as WHU in study). The structure of spike glycoprotein of delta and omicron variant was determined using spike-WHU (pdb ID: 7DF4) as template in Modeller v9.3. 100 structures were generated for each variant and final structure was selected on the basis of DOPE score and Ramachandran plot assessment.

In order to identify the interacting residues of spike-delta and spike-omicron with ACE2, docking was performed through HADDOCK. The binding energies were calculated using PRODIGY. The interaction profile of spike-ACE2-delta and spike-ACE2-omicron complex were further explored through pdbSUM and PyMol.

### Electrostatic potential

The electrostatic potential of SARS-CoV-2 spike protein of WHU, delta and omicron was calculated through Delphi and PyMOL.

## Results

### Sequence Analysis

The sequence alignment of spike protein of delta and omicron variants with WHU01 confirm the presence of mutations in the spike gene. The important mutations of omicron identified in the receptor binding domain are G339D, S371L, S373P, S375F, S477N, T478K, E484A, Q493R, G496S, Q498R, N501Y, and Y505H. The significant mutation in the furin cleavage site is P681H. Whereas in case of delta variant only two mutations (L452R, T478K) were observed in the receptor binding domain.

### Interaction Analysis

The interaction analysis of spike-ACE2-WHU revealed that important residues of spike-WHU at the receptor binding interface were F486, N487, N501, Y505, G502, T500, Q493 and S494. In total 18 residues of ACE-2 and 19 residues of spike-WHU were present at the interface. The important H-bonding interactions were observed as S494:H34, Q493:E35, and G502:K353. Q493 also involved in forming salt bridge with K31 (Table 1).

**Table 1:** Interactions of spike-WHU (pdb ID: 7DF4) with ACE2.

| S. No | Spike-WHU | ACE-2 | Distance (Å) |
| --- | --- | --- | --- |
| 1. | Phe-486 | Met-82 | 3.77 |
| 2. | Asn-487 | Tyr-83 | 3.8 |
| 3. | Gln-493 | Lys-31<br>Glu-35 | 3.7<br>2.9 |
| 4. | Ser-494 | His-34 | 2.7 |
| 5. | Tyr-505 | Asp-38 | 3.5 |
| 6. | Gly-502 | Lys-353 | 2.4 |
| 7. | Asn-501 | Lys-353 | 3.2 |
| 8. | Thr-500 | Tyr-41<br>Asp-355 | 3.2<br>3.4 |

The interaction pattern of delta variant revealed the interacting residues as F486, N487, N501, G502, T500, K478, R403, Y505, Q493 and S494. There were 18 residues of ACE2 and 21 residues of delta-spike protein at the interface that involved in interactions. The important H-bonding interactions observed as S494:H34, Y505:D38. The T478K mutation resulted in the involvement of 478K in interactions with ACE2 as K478:Q24 (Table2). The T478K mutation in delta variant may have an impact on increasing the binding affinity with ACE2 due to addition of a charged amino acid (K) in place of an aliphatic amino acid (T).

**Table 2:** Interactions of spike-delta with ACE2.

| S. No | Spike-Delta | ACE-2 | Distance (Å) |
| --- | --- | --- | --- |
| 1. | Phe-486 | Met-82 | 3.6 |
| 2. | Asn-487 | Tyr-83 | 3.4 |
| 3. | Gln-493 | Lys-31<br>Glu-35 | 3.2<br>3.5 |
| 4. | Ser-494 | His-34 | 2.4 |
| 5. | Tyr-505 | Asp-38 | 2.5 |
| 6. | Gly-502 | Lys-353 | 2.7 |
| 7. | Asn-501 | Lys-353 | 2.4 |
| 8. | Thr-500 | Tyr-41<br>Asp-355 | 3.0<br>2.8 |
| 9. | Lys-478 | Gln-24 | 3.1 |
| 10. | Arg-403 | His-34 | 3.2 |

There were 26 residues of Spike and 22 residues of ACE2 at the interface that involved in interactions. The important H-bonding interactions observed are S494:H34, S496:D38, R498:Y41, Y501:K353, and H505:R393. The salt bridge observed as R493:D38, and H505:E37. With the mutations, a large number of interaction have been observed in the residues lying in the domain of 490-505 (Table 3). The Q493R and Q498R mutations may result in the strong binding interactions due to the replacement of polar charged amino acid. The replacement of polar residue in place of hydrophobic residue at G496S may result in the strong binding interactions. The Y505H mutation result in replacement of polar aliphatic amino acid with charged amino acid which can further increased binding interactions. The involvement of polar and charged amino acids in interactions with the ACE2 can not only result in strong binding but may also increase the number of interactions between spike-omicron and ACE2.

**Table 3:**
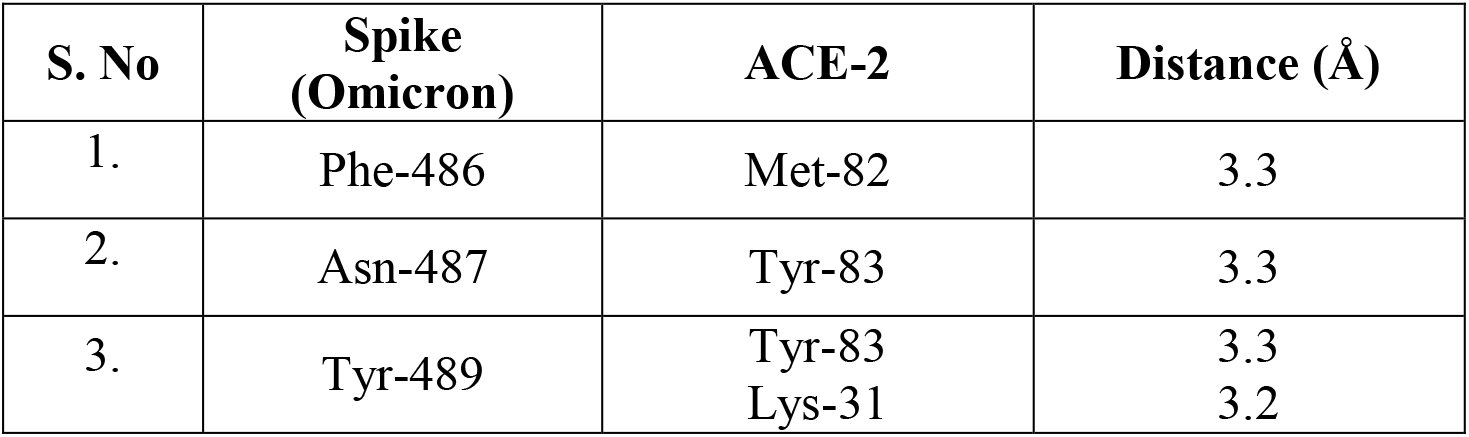
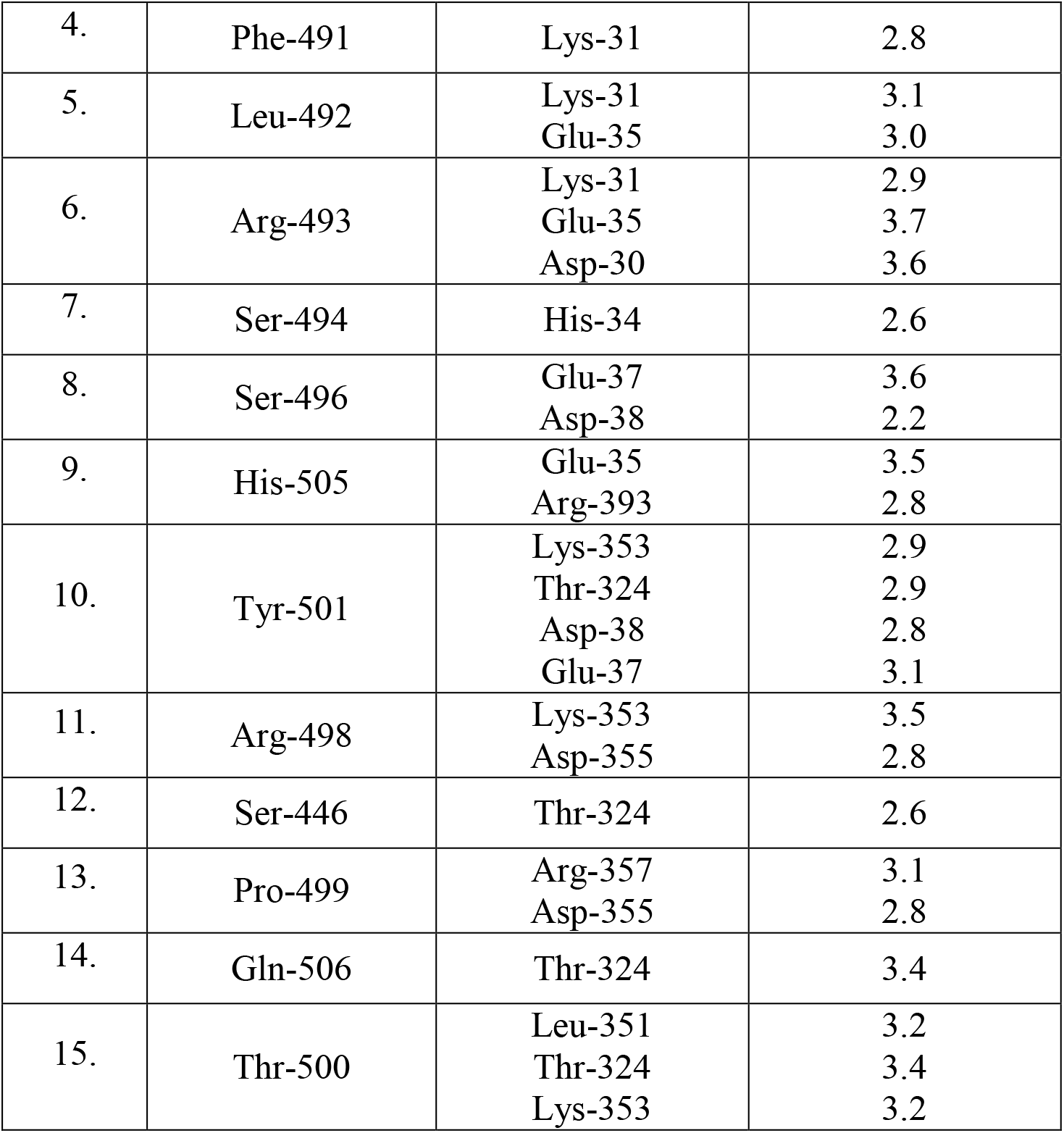
Interactions of spike-omicron with ACE2.

| S. No | Spike (Omicron) | ACE-2 | Distance (Å) |
| --- | --- | --- | --- |
| 1. | Phe-486 | Met-82 | 3.3 |
| 2. | Asn-487 | Tyr-83 | 3.3 |
| 3. | Tyr-489 | Tyr-83<br>Lys-31 | 3.3<br>3.2 |
| 4. | Phe-491 | Lys-31 | 2.8 |
| 5. | Leu-492 | Lys-31<br>Glu-35 | 3.1<br>3.0 |
| 6. | Arg-493 | Lys-31<br>Glu-35<br>Asp-30 | 2.9<br>3.7<br>3.6 |
| 7. | Ser-494 | His-34 | 2.6 |
| 8. | Ser-496 | Glu-37<br>Asp-38 | 3.6<br>2.2 |
| 9. | His-505 | Glu-35<br>Arg-393 | 3.5<br>2.8 |
| 10. | Tyr-501 | Lys-353<br>Thr-324<br>Asp-38<br>Glu-37 | 2.9<br>2.9<br>2.8<br>3.1 |
| 11. | Arg-498 | Lys-353<br>Asp-355 | 3.5<br>2.8 |
| 12. | Ser-446 | Thr-324 | 2.6 |
| 13. | Pro-499 | Arg-357<br>Asp-355 | 3.1<br>2.8 |
| 14. | Gln-506 | Thr-324 | 3.4 |
| 15. | Thr-500 | Leu-351<br>Thr-324<br>Lys-353 | 3.2<br>3.4<br>3.2 |

### Comparison of Interactions

The Table 4 summarizes the comparison of interactions among spike-WHU, -delta and -omicron complexes with ACE2. In terms of binding energy, it has been observed that spike-ACE2-omicron complex was having the highest binding affinity of −17.6Kcal/mol compared to spike-ACE2-WHU with −13.2Kcal/mol and spike-ACE2-delta with −13.3Kcal/mol. In terms of dissociation constant (Kd), the spike-ACE2-omicron complex was having the lowest dissociation constant (1.3E-13 M) hence having high binding affinity. When comparing the residues at the binding interface, the spike-ACE2-omicorn complex was having highest number of charged and polar residues at the interface compared to spike-ACE2-WHU and spike-ACE2-delta complex (Table 5). While comparing the interaction pattern, in spike-ACE2-omicorn complex the new interactions were observed with the residues of spike protein as Y489, F491, L492, S496, R498, Q506, P499, and S446. All of these residues were not involved in interactions in spike-ACE2-WHU complex as well as spike-ACE2-delta complex. The large number of mutations in the receptor binding domain including Q493R, N501Y, Y505H, and T478K may result in the conformational changes that lead to development of new interactions in the receptor binding domain.

**Table 4:** Comparison of interactions of wild-ACE2, delta-ACE2, and Omicron-ACE2.

|  | WHU | Delta | Omicron |
| --- | --- | --- | --- |
| **Binding Affinity**<br>**$\Delta G$ (Kcal/mol)** | -13.2 | -13.3 | -17.6 |
| **Dissociation**<br>**Constant $K_d$ (M)** | 2.2E-10 | 1.9E-10 | 1.3E-13 |
| **Number of Interfacial Contacts (ICs) per property** |  |  |  |
| **charged-charged** | 5 | 5 | 18 |
| **charged-polar** | 10 | 13 | 20 |
| **charged-apolar** | 23 | 22 | 49 |
| **polar-polar** | 5 | 7 | 6 |
| **polar-apolar** | 26 | 29 | 30 |
| **apolar-apolar** | 13 | 16 | 31 |

### Electrostatic Potential of Spike Glycoprotein

The electrostatic surface characteristics also revealed that spike-WHU receptor binding domain is less electropositive with electro positivity slightly increased in case of spike-delta while spike-omicron is more electropositive. This can be explained by the mutations observed in the receptor binding domain of omicron as S477N, T478K, Q493R, G496S, Q498R, and Y505H.

## Discussion

The Omicron variant of SARS-CoV-2 has emerged with significant number of mutations in the spike gene some of which are novel as G339D, S371L, S373P, S375F, N440K, G446S, E484A, Q493R, G496S, Q498R, Y505H, T547K, H355Y, N679K, N764K, D796Y, N856K, Q954H, N969K, and L981F. The new mutations result in the addition of polar and charged residues in the spike protein as Arg, Lys, His, and Asp. The addition of polar and charged residues may have an impact on increasing the receptor binding interactions of spike protein with ACE2.

In the current study the interaction analysis of spike-omicron has revealed that this variant has strong interactions with the ACE2 receptor compared to WHU strain and delta variant. The spike-ACE2-delta complex has almost similar binding affinity with spike-ACE2-WHU complex. It has been shown in a study that the delta variant with K417N, L452R, and T478K mutations have similar binding affinity for ACE2 as observed in wild type spike protein. But in case of the N501Y mutation the affinity has increased [4]. The increased binding affinity of spike-omicron with ACE2 could be explained by the addition of large number of charged and polar residues in the spike protein particularly in the receptor binding domain. The strong binding affinity of spike-omicron with ACE2 may result in increased transmissibility and infectivity of the variant. This can be explained by looking at the spread of virus, since the identification of virus in South Africa it has spread to 41 countries around the globe with large number of cases observed in South Africa, United Kingdom, USA, and Australia.

The study results have shown an apparent increase in binding affinity for ACE2 receptor as a result of mutations observed in Omicron variant. The replacement of aliphatic residues with polar charged residues with resulting increased electropositivity at crucial RBD residues appear to be the attributable reason, however *in-vivo* cell line studies can suggest a more confirmatory mark on the said findings.

**Figure 1:**
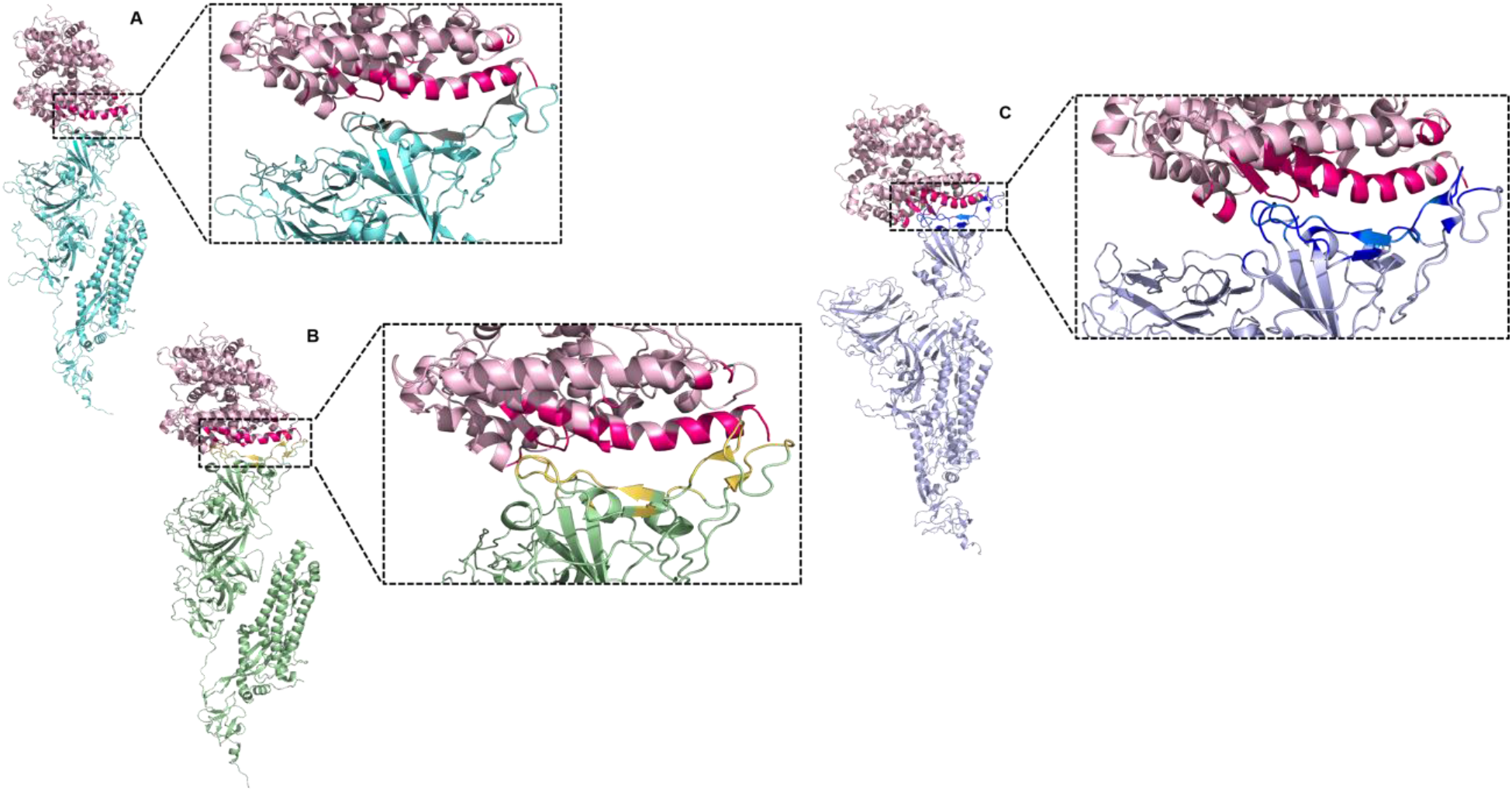
Structure of sARS-CoV-2 spike glycoprotein in complex with ACE2. (A) The spike-ACE2-WHU complex. The spike protein is represented in torquoise color. (B) spike-ACE2-delta complex. The spike-delta is represented in green color and the binding interface is shown in yellow color. (C) Spike-ACE2-omicron complex. The spike-omicron is represented in silver color with binding interface highlighted as blue. The ACE2 protein is shown in pink color.

**Figure 2:**
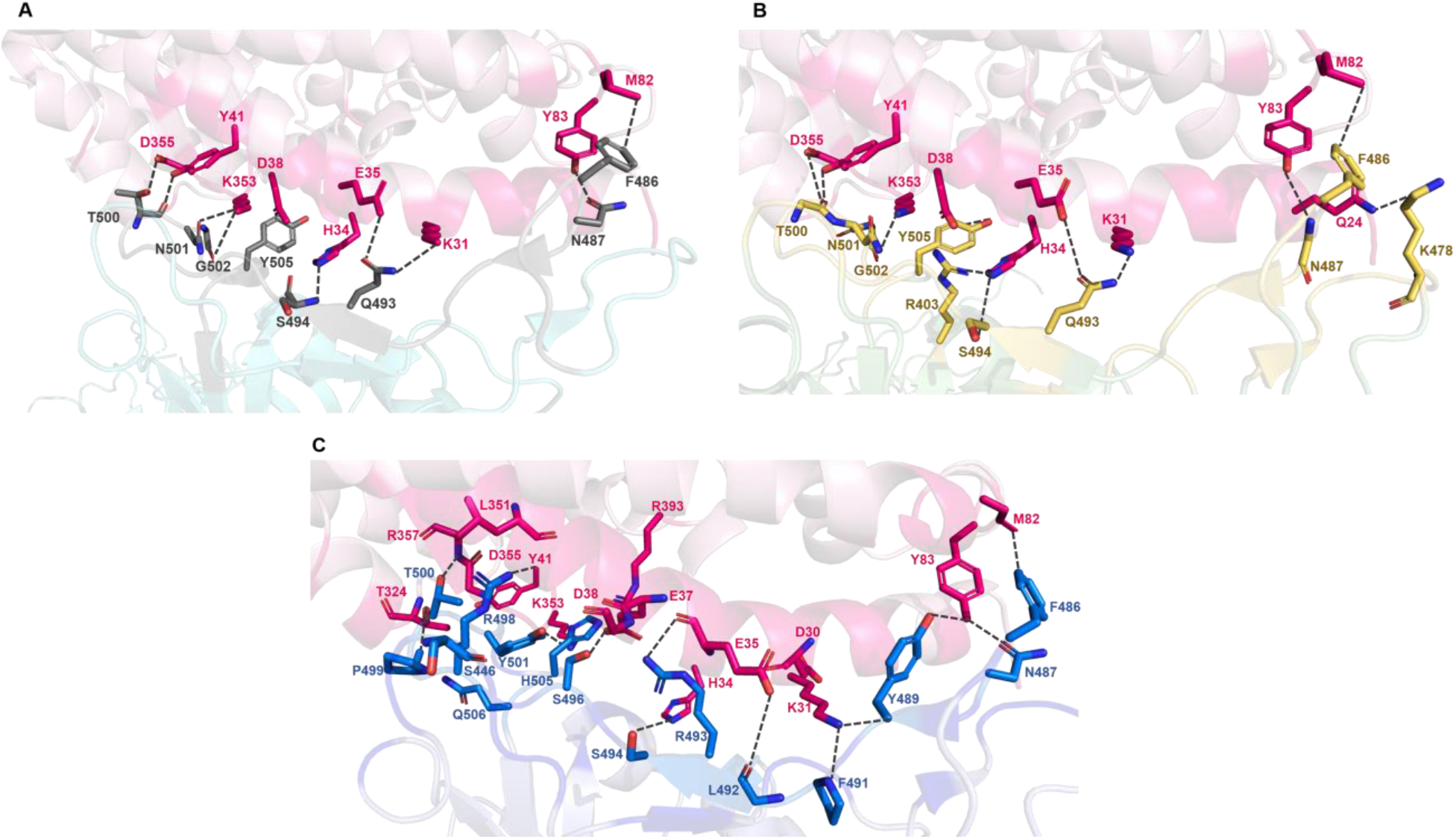
The interactions of ACE2 with WHU-spike, delta-spike, and omicron-spike. (A) spike-ACE2-WHU complex. The interacting residues of WHU-spike protein are show in grey stick representation. (B) Spike-ACE2-delta complex. The interacting residues of delta-spike are shown in yellow stick representation. (C) spike-ACE2-omicron complex. The interacting residues of delta-spike are shown in blue stick representation.

**Figure 3:**
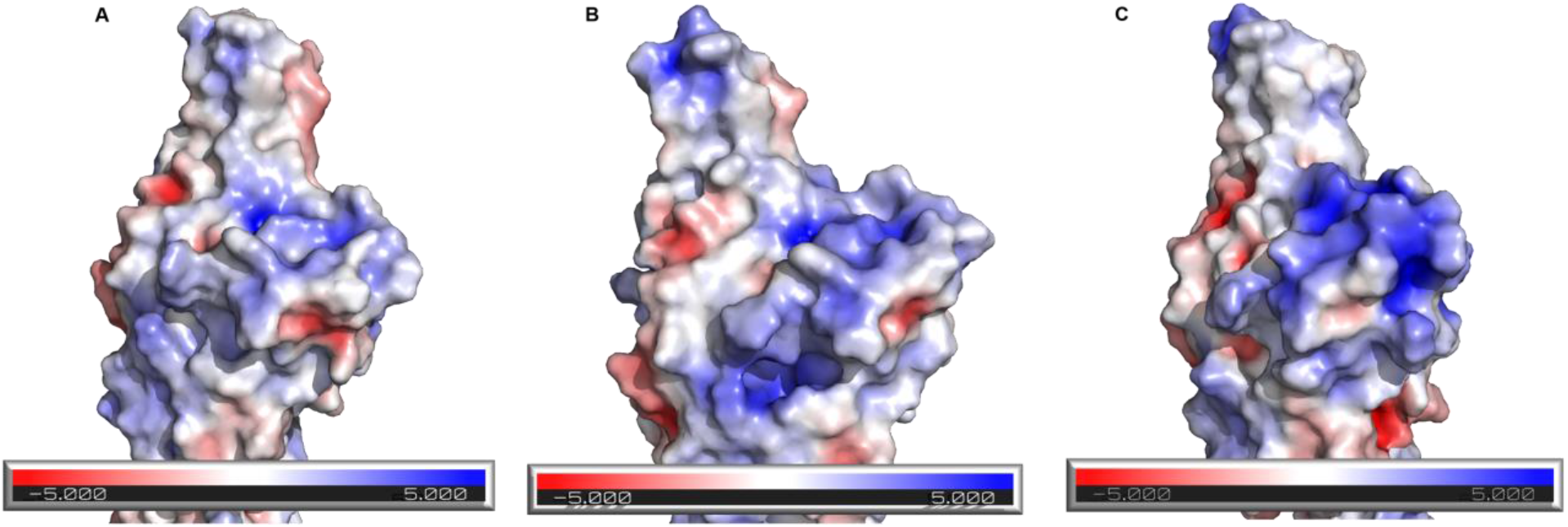
Electrostatic potential of receptor binding domain of spike protein. (A) spike-WHU; (B) spike-delta; (C) spike-omicron. The positive charge is represented by blue color while negative charge is represented by red color. The electropositive potential of receptor binding domain gradually increases from spike-WHU to spike-omicron.

